# Prevention of EloR/KhpA heterodimerization by introduction of site-specific amino acid substitutions renders the essential elongasome protein PBP2b redundant in *Streptococcus pneumoniae*

**DOI:** 10.1101/435529

**Authors:** Anja Ruud Winther, Morten Kjos, Gro Anita Stamsås, Leiv Sigve Håvarstein, Daniel Straume

## Abstract

The RNA binding proteins EloR and KhpA are important components of the regulatory network that controls and coordinates cell elongation and division in *S. pneumoniae*. Loss of either protein reduce cell length, and makes the essential elongasome proteins PBP2b and RodA dispensable. It has been shown previously in formaldehyde crosslinking experiments that EloR co-precipitates with KhpA, indicating that they form a complex *in vivo*. In the present study, we used 3D modeling and site directed mutagenesis in combination with protein crosslinking to further study the relationship between EloR and KhpA. Protein-protein interaction studies demonstrated that KhpA forms homodimers and that KhpA in addition binds strongly to the KH-II domain of EloR. Site directed mutagenesis identified isoleucine 61 (I61) as crucial for KhpA homodimerization. When substituting I61 with phenylalanine, KhpA lost the ability to homodimerize, while it still interacted strongly with EloR. In contrast, both homo- and heterodimerization were lost when I61 was substituted with tyrosine. By expressing these KhpA versions in *S. pneumoniae*, we were able to show that disruption of EloR/KhpA heterodimerization makes the elongasome redundant in *S. pneumoniae*. Of note, loss of KhpA homodimerization did not give rise to this phenotype, demonstrating that the EloR/KhpA complex is crucial for regulating the activity of the elongasome. In support of this conclusion, we found that localization of KhpA to the pneumococcal mid-cell region depends on its interaction with EloR. Furthermore, we found that the EloR/KhpA complex co-localizes with FtsZ throughout the cell cycle.

**Importance:** To ensure correct cell division, bacteria need to monitor the progression of cell division and coordinate the activities of cell division proteins accordingly. Understanding the molecular mechanisms behind these regulatory systems is of high academic interest and might facilitate the development of new therapeutics and strategies to combat pathogens. EloR and KhpA form a heterodimer that is part of a signaling pathway controlling cell elongation in the human pathogen *S. pneumoniae*. Here we have identified amino acids that are crucial for EloR/KhpA heterodimerization, and demonstrated that disruption of the EloR/KhpA interaction renders the cells independent of a functional elongasome. Furthermore, we found the EloR/KhpA complex to co-localize with the division ring (FtsZ) during cell division.

## Introduction

In most bacteria, the cytoplasmic membrane is surrounded by a peptidoglycan layer, which gives the cell its shape and provides resistance to internal turgor pressure (1). The peptidoglycan sacculus also serves as an anchoring device for surface proteins and other cell wall components such as teichoic acids and extracellular polysaccharides (2–5). During cell division and growth, the peptidoglycan synthesis machineries add new material into the existing cell wall. In ovoid bacteria, such as the important human pathogen *Streptococcus pneumoniae*, two modes of cell wall synthesis occur. The divisome synthesizes the septal crosswall, while extension of the lateral cell body is carried out by the elongasome (6, 7). The cell wall synthesis machineries of *S. pneumoniae* contain six penicillin binding proteins (PBPs), five of which participate in building the cell wall via transglycosylase and transpeptidase reactions. The class A PBPs, PBP1a, PBP2a, PBP1b, perform both reactions, while the class B PBPs, PBP2b and PBP2x, only have transpeptidase activity. Recently, it was discovered that the monofunctional class B enzymes PBP2x and PBP2b operate in conjunction with FtsW and RodA, two newly discovered transglycosylases belonging to the SEDS family proteins (<u>s</u>hape, <u>e</u>longation, <u>d</u>ivision and <u>s</u>porulation) (8, 9). The sixth PBP, PBP3, is a D,D-carboxypeptidase that reduces the level of inter peptide cross-bridges in the peptidoglycan by cleaving off the C-terminal D-Ala residue in stem pentapeptides (10). PBP2b and RodA have been found to be essential for cell elongation, while PBP2x and FtsW are essential for synthesis of the septal disc. Functional studies and subcellular localizations suggest that PBP2b/RodA and PBP2x/FtsW are key components of the elongasome and the divisome, respectively (11–14). It is not clear whether the elongasome- and divisome activities alternate or if these machineries work simultaneously during cell division. However, some data suggest a short period of cell elongation before the onset of septal peptidoglycan synthesis (12, 15).

In contrast to rod-shaped bacteria, *S. pneumoniae* lacks MreB, a cytoskeleton-like protein that moves the cell wall synthesis machinery in helical patterns perpendicular to the cell length (16). Instead, pneumococci elongate by inserting new peptidoglycan into the existing cell wall between the future cell equator and the septum in a circumferentially motion guided by the FtsZ/FtsA division ring (6, 17, 18). At some point during cell elongation, the divisome initiates septal cross wall synthesis. If the coordinated activities of the elongasome and the divisome get out of control, it leads to severe growth defects and development of morphological abnormalities (11, 13, 19). The cells have therefore developed sophisticated systems to monitor cell cycle progression in order to fine-tune the activity of the elongasome and divisome during cell division. One of these systems includes the membrane-spanning eukaryotic-like serine/threonine kinase StkP. It has four extracellular cell-wall-binding PASTA domains, which are believed to monitor the status of the cell wall during division and activate the appropriate cell division proteins through phosphorylation (20–23).

In a recent study we found that EloR, which is phosphorylated by StkP on threonine 89 (24, 25), is a key regulator of cell elongation in *S. pneumoniae* (26). Our results indicated that EloR stimulates cell elongation when phosphorylated, while being inactive or preventing elongation in its non-phosphorylated form. Moreover, we found that Δ*eloR* cells can survive without PBP2b and its cognate SEDS transglycosylase RodA, demonstrating that deletion of *eloR* supresses the need for a functional elongasome in *S. pneumoniae*. Cells lacking EloR displayed a significant reduction in growth rate and became short and round (25, 26). EloR is a cytoplasmic protein of 37 kDa comprising three different domains: an N-terminal jag-domain of unknown function followed by two RNA-binding domains, a type II KH domain (KH-II) and R3H, at the C-terminal end (27, 28). In a recent study Zheng et al. (29) showed that EloR co-precipitates with a protein called KhpA after treating cells with formaldehyde cross linker. KhpA is a small (8.9 kDa) RNA-binding protein that consists only of a type II KH domain. Similar to EloR, deletion of the *khpA* gene supresses the need for a fully functional elongasome, as *pbp2b* as well as *rodA* can be deleted in a Δ*khpA* mutant (29). EloR and KhpA probably bind certain target RNAs to modulate expression of specific cell division and/or elongation proteins during different stages of the cell cycle. In support of this hypothesis Zheng et al. (29) reported that the absence of EloR or KhpA results in higher cellular levels of the cell division protein FtsA, and that this increase compensates for the loss of PBP2b (29). Homologs of EloR and KhpA appear to be widespread in many Gram-positive bacteria, and are found in genera such as *Streptococcus*, *Bacillus*, *Clostridium*, *Listeria*, *Enterococcus*, *Lactobacillus* and *Lactococcus*. The conservation of these proteins across large phylogenetic distances indicates that they are central players in the cell elongation and division machineries of low G+C Gram-positive bacteria.

In the present study, we show that KhpA homodimerizes, and that it in addition interacts strongly with the KH-II domain of EloR forming an EloR/KhpA heterodimer. Furthermore, we identified amino acids critical for these interactions. We successfully constructed a single amino acid mutant of KhpA that fails to homodimerize but still interacts with EloR, and a single amino acid mutant that neither self-interacts nor heterodimerizes. The unique properties of these KhpA versions were used to demonstrate that the function of EloR is compromised when it is no longer able to interact with KhpA, resulting in cells phenocopying Δ*eloR* and Δ*khpA* mutants (reduced cell elongation). Finally, *in vivo* localization studies showed that KhpA co-localizes with FtsZ throughout the cell cycle, and that this localization pattern depends on its interaction with EloR.

## Results

### KhpA interacts with itself and the KH-II-domain of EloR

In a recent study we showed that the loss of EloR suppresses the need of a functional elongasome in *S. pneumoniae* since *pbp2b* and *rodA* could be deleted (26). Soon after this, Zheng and co-workers published that EloR co-precipitated with a small protein (8.9 kDa) called KhpA in formaldehyde crosslinking experiments. In addition, they found that a Δ*khpA* mutant phenocopies a Δ*eloR* mutant and that both proteins bound to a similar set of RNA molecules in pulldown experiments (29). In the present work, we utilized a bacterial two-hybrid system (BACTH assay) to further study the interaction between EloR and KhpA. The BACTH system is based on interaction-mediated reconstitution of the *Bordetella pertussis* adenylate cyclase CyaA, which consists of two domains (T18 or T25). When brought together through interaction of the proteins tested, the active T18-T25 reconstitution produces cAMP, which ultimately results in measurable β-galactosidase production in the *E. coli* host (30). When testing full-length EloR against KhpA in the BACTH assay, we observed a strong positive interaction (Fig. 1), confirming the crosslinking results of Zheng and co-workers (29). Next, we wanted to identify the part of EloR that interacts with KhpA. To do so, each of the three domains of EloR (Jag, KH-II and R3H) was tested individually against KhpA (Fig. 1). The results clearly showed that KhpA specifically interacts with the KH-II-domain of EloR (KH-II^EloR^).

**Fig. 1.**
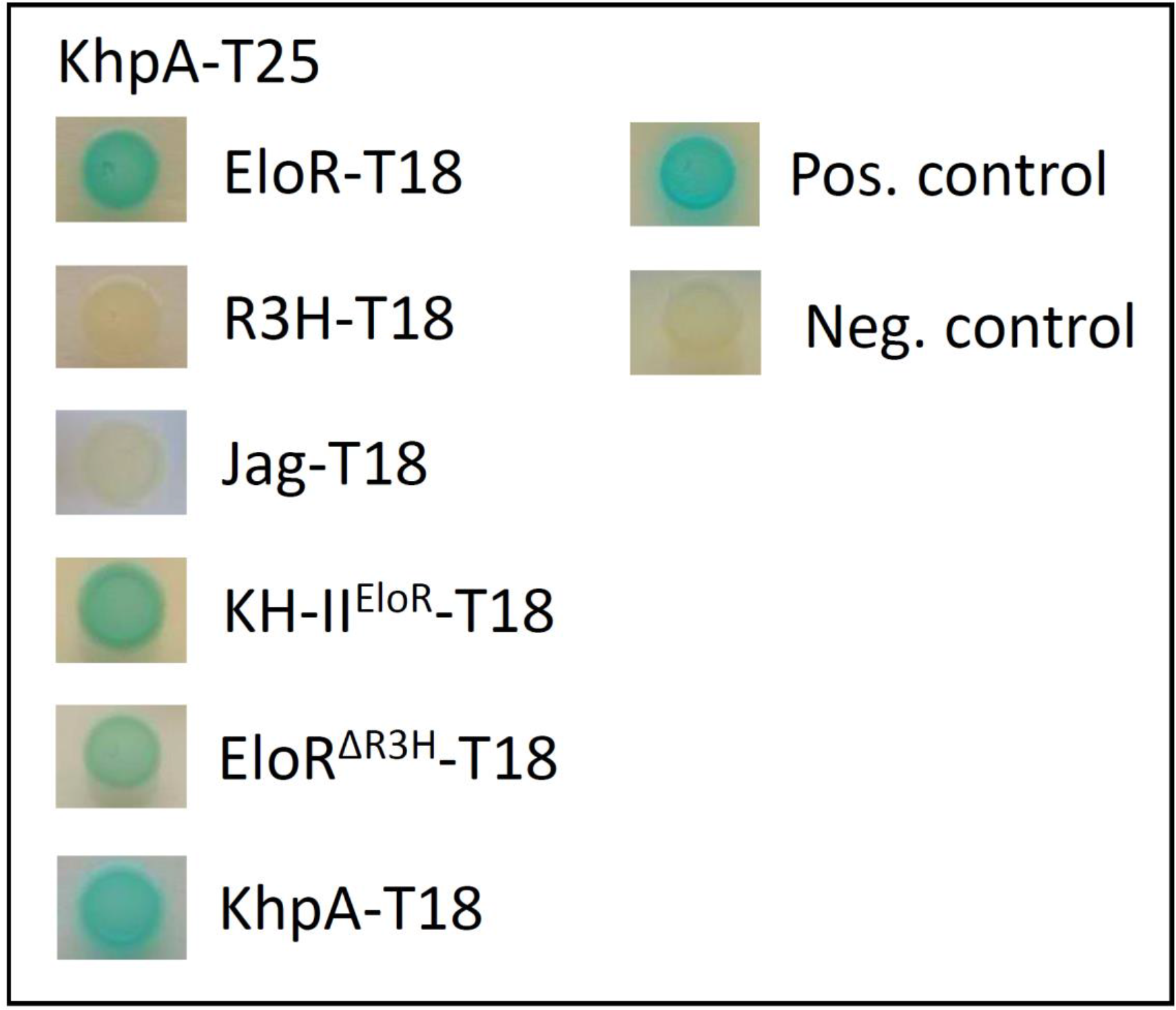
BACTH-assay showing that KhpA interacts directly with EloR and with itself. KhpA was probed against full-length EloR, the R3H domain, the KH-II^EloR^ domain, the Jag domain and EloR missing the C-terminal R3H domain (EloR^ΔR3H^). Positive interactions (blue spots) were only seen between KhpA and parts of EloR having the KH-II^EloR^ domain. The positive self-interaction of KhpA is shown at the bottom.

Since KH-domains recognize on average up to four nucleotides, they have a tendency to interact with each other to bind longer sequences and thereby increase their target specificity (28, 31). We therefore suspected that KhpA self-interacts and forms homodimers. BACTH assays using KhpA fused to T18 and T25 resulted in a strong positive signal (Fig. 1), suggesting that KhpA, in addition to interacting with EloR, also forms homodimers.

### Identification of amino acid residues crucial for KhpA homo- and EloR/KhpA heterodimerization

We reasoned that a 3D model of KhpA might help us identify amino acids that are crucial for homodimerization and heterodimerization with EloR. KH-domains have a highly conserved fold and many 3D-structures are available in the databases (28, 31). To predict the 3D structure of KhpA, we used the online structure prediction tool iTasser. As expected, the predicted structure shows a typical KH-II domain (C-score = −0.36) consisting of three α-helices packed against a three-stranded β-sheet (α-β-β-α-α-β) (Fig. 2A). The conserved RNA binding cleft is made up of the third α-helix and the third β-strand. The typical GxxG loop that interacts with the phosphate backbone of the ssRNA (or in some cases ssDNA) is located between the α2- and α3-helices (marked in green in Fig. 2A). Introduction of two aspartates in this loop (GDDG) abolishes binding of target RNA (32). To predict the interaction surface between two KhpA molecules, we did protein docking using ZDOCK with the 3D-model of KhpA as input. According to the model (ZDOCK score = 895.421), the α3-helix creates an anti-parallel interaction surface between two KhpA proteins, resulting in a homodimeric structure where the GxxG loops of the two proteins point in opposite directions (Fig. 2B). Based on this structure, we made four different mutant versions of KhpA in which single amino acids predicted to protrude from the α3-helix was altered (R53K, R59K, T60Q and I61F). The point mutated versions of KhpA where then tested for their ability to homodimerize by performing BACTH assays. The changes in position 53, 59 or 60 did not dramatically reduce homodimerization, but changing I61 to the bulkier phenylalanine abolished the interaction between KhpA monomers (Fig. 2C). To get more accurate data on the effect of the I61F mutation, quantitative measurements of the β-galactosidase production were performed (see Materials and Methods). Indeed, the KhpA^I61F^ mutant protein has completely lost the ability to self-interact, but can still form heterodimers with EloR (Fig. 3A). In an attempt to create a KhpA mutant that does not form homodimers nor EloR/KhpA heterodimers, I61 was changed to tyrosine, which adds a polar hydroxyl group to the bulky phenyl ring. When tested in quantitative BACTH assays, our results showed that the KhpA^I61Y^ mutant has lost the ability to interact with itself and the interaction with EloR was dramatically reduced (Fig. 3A).

**Fig. 2.**
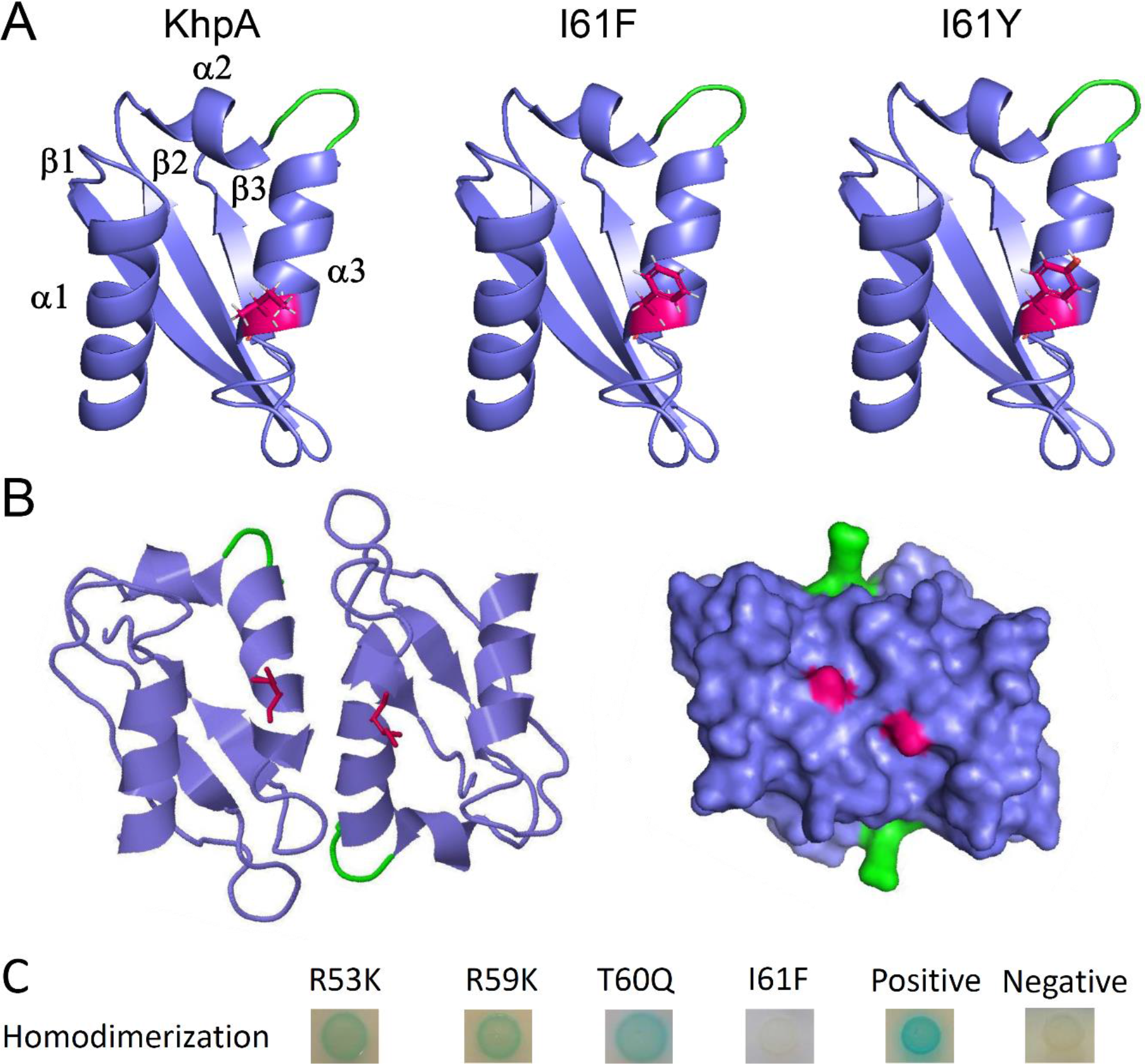
Structure prediction of KhpA using iTasser and ZDOCK. A. KhpA was predicted to have the typical α-β-β-α-α-β fold of KH-II domains, with the I61 (shown in magenta) protruding from the α3-helix. The structures of the I61F and I61Y substitutions are shown. B. Protein-protein docking of KhpA homodimers using ZDOCK. The α3-helix of two KhpA molecules are predicted to make contact anti-parallel of each other forming a homodimer where the GXXG RNA-binding loops (shown in green) point in opposite directions. The I61 (magenta) of two KhpA monomers are brought in close proximity in the dimeric structure, facilitating a hydrophobic contact surface. C. BACTH assay showing KhpA’s ability to form homodimers when selected amino acids in the α3-helix were changed (R53K, R59K, T60Q and I61F). Positive interactions appear as blue spots.

**Fig. 3.**
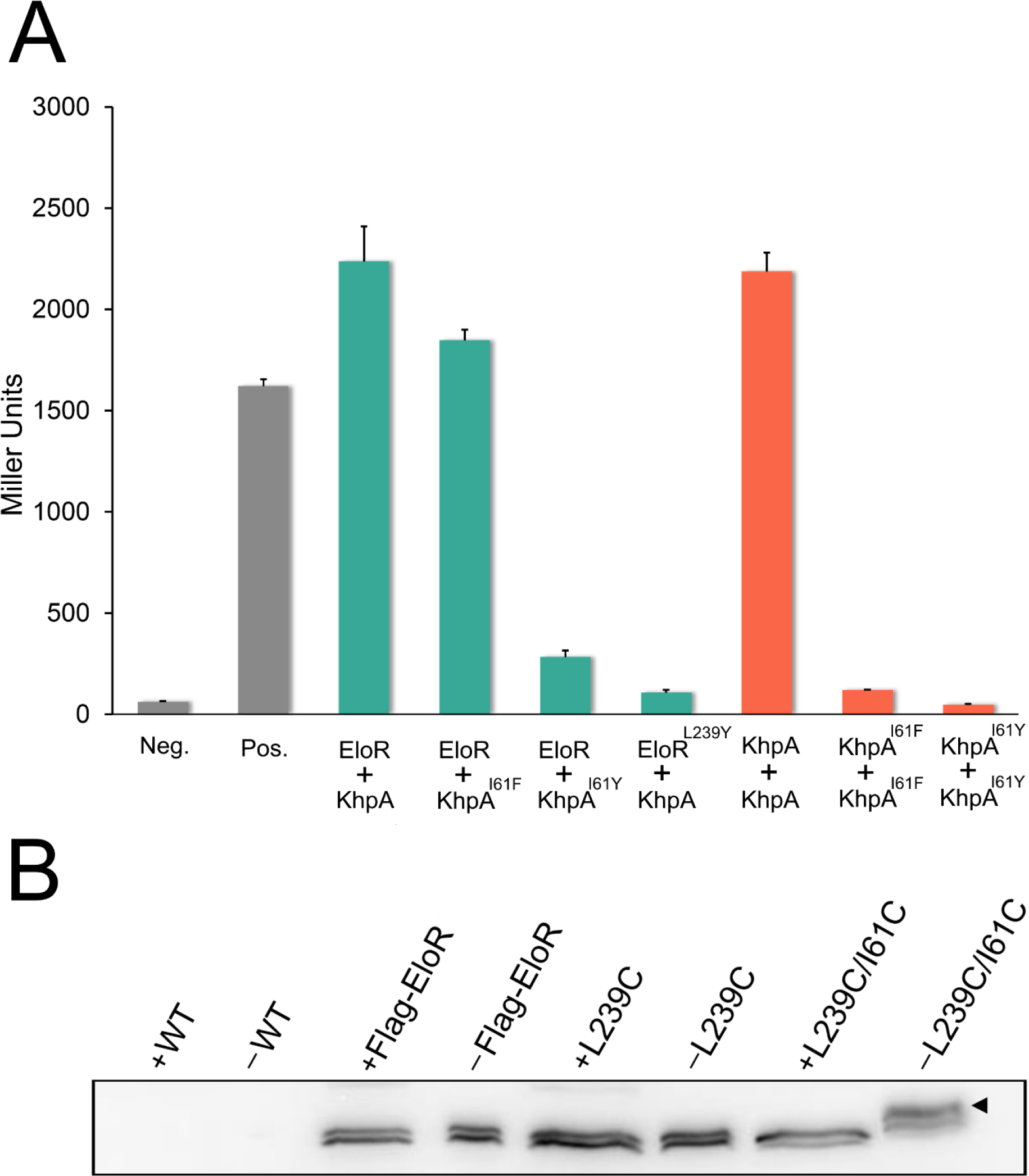
The α3-helix of KhpA is critical for self-dimerization and for EloR/KhpA complex formation. A. Quantitative measurements of β-galactosidase production in BACTH assays testing the interaction between EloR and KhpA, KhpA^I61F^ or KhpA^I61Y^ in addition to EloR^L239Y^ against KhpA (green bars). β-galactosidase production resulting from homodimerization of KhpA, KhpA^I61F^ and KhpA^I61Y^ is represented by orange bars, while negative and positive controls are shown in grey. B. Immunoblot detection of 3xflag-EloR in strain RH425, SPH448, AW334 and AW336. A crosslinked EloR/KhpA complex was observed in strain AW336 under non-reducing conditions (-), but not after reduction with β-mercaptoethanol (+).

Amino acid sequence alignment of the KH-II^EloR^ and KhpA, suggests that leucine 239 (L239) in EloR corresponds to I61 in KhpA (see supplemental Fig. S1). Accordingly, when L239 in EloR was substituted with a tyrosine, KhpA could no longer interact with EloR, showing that this residue is indeed important for EloR/KhpA heterodimerization (Fig. 3A). To prove that L239 and I61 are in close proximity in the EloR/KhpA heterodimer, we replaced these two amino acids with cysteins to determine whether this would result in a disulfide bridge between the two proteins *in vivo*. A pneumococcal strain expressing the mutant proteins EloR^L239C^ and KhpA^I61C^ was therefore constructed (strain AW336). EloR^L239C^ contained an N-terminal 3xflag-tag to allow detection with α-flag antibodies. Strain AW336 was grown to exponential phase, harvested, and lysed using SDS loading buffer with or without the reducing agent β-mercaptoethanol (see Material and Methods). Next, samples were analyzed by SDS-PAGE followed by immunoblotting. In non-reduced cell lysates, we detected a shift in band size corresponding to the complex between EloR and KhpA (Fig. 3B). This shift was not present in samples where β-mercaptoethanol had been added to break the disulfide bond, or in any of the samples containing wild type 3xflag-EloR or 3xflag-EloR^L239C^ only. This confirms the interaction between KhpA and the KH-II domain of EloR *in vivo*, and that I61 in the α3-helix of KhpA interacts directly with L239 in the α3-helix of the KH-II^EloR^ domain.

### Prevention of EloR/KhpA heterodimerization relieves the requirement of *pbp2b*

A Δ*khpA* mutant phenocopies a Δ*eloR* mutant (29). Both mutants have reduced growth rates, form shorter cells and are viable without a functional elongasome (i.e. without a *pbp2b* or *rodA* gene) (26, 29). We hypothesized that the reason Δ*khpA* cells phenocopies Δ*eloR* cells is because deletion of either will prevent the formation of the EloR/KhpA complex. In other words, the elongasome only becomes essential when the EloR/KhpA complex is able to form and carry out its normal biological function. To test this hypothesis we exploited the unique properties of KhpA^I61F^ and KhpA^I61Y^. KhpA^I61F^ does not form homodimers, but form heterodimers with EloR, while KhpA^I61Y^ is unable to form either. First, we examined if expression of KhpA^I61F^ or KhpA^I61Y^ generated cells with reduced growth rate similar to a Δ*kphA* mutant. Deletion of *khpA* (strain DS420) increased the doubling time with approximately 15 minutes, which complies with previous findings (15-30 minutes) (29), while strains expressing KhpA^I61F^ or KhpA^I61Y^ (AW212 and AW275) had growth rates similar to the wild type strain (data not shown). Microscopic examination of KhpA^I61F^ or KhpA^I61Y^ cells showed that the KhpA^I61Y^ strain grew in short chains similar to KhpA deficient cells. The KhpA^I61F^ strain on the other hand grew mainly as diplococci similar to the wild type strain (Fig. 4A). By measuring cell lengths and widths, it became evident that KhpA^I61Y^ cells, in which KhpA is unable to form a complex with EloR, have a rounder cell morphology with reduced cell elongation similar to Δ*khpA* cells (Fig. 4B). This phenotype is also characteristic for Δ*eloR* cells (25, 26, 29). In contrast, cells expressing the monomeric version of KhpA (I61F) that can still form a complex with EloR, displayed a normal length/width distribution (Fig. 4B).

**Fig. 4.**
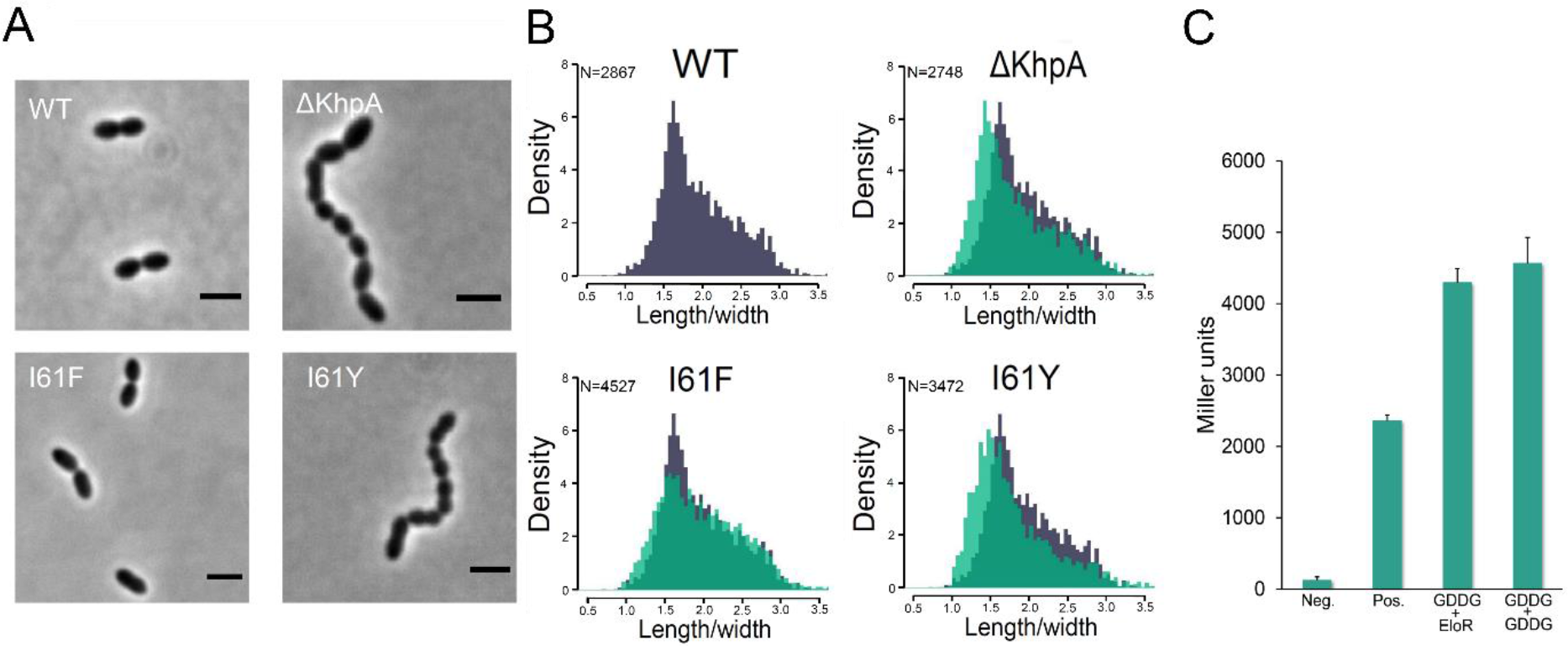
A. Comparison of the morphology of strain RH425 (wt), DS420 (Δ*khpA*), AW212 (I61F) and AW275 (I61Y). Loss of KhpA homodimerization (KhpA^I61F^) produced cells with morphology similar to wild type. Cells in which KhpA no longer interacts with EloR (KhpA^I61Y^) had morphologies resembling the Δ*khpA* mutant. Scale bars are 2 μm. B. Comparison of the cell-shape distribution (length/width) of Δ*khpA*-, KhpA^I61F^- and KhpA^I61Y^- cells (in green) with wild type cells (in grey). KhpA^I61Y^ and Δ*khpA* cells were both significantly different from wild type (p<0.05, two-sample t-test), while the shape distribution of KhpA^I61F^ cells was similar to wild type. C. Quantitative BACTH assay showing that KhpA^GDDG^ self-dimerizes and forms complex with EloR.

To further test our hypothesis that EloR/KhpA heterodimerization is required for normal elongasome function, we compared pneumococcal mutants expressing KhpA^I61F^, KhpA^I61Y^ and EloR^L239Y^ (AW279) with respect to the essentiality of their *pbp2b* gene. Indeed, *pbp2b* could be deleted in KhpA^I61Y^ and EloR^L239Y^ cells with normal transformation frequencies, but not in KhpA^I61F^ cells. Since it has been shown that mutants expressing a KhpA unable to bind ssRNA (changing the ssRNA-binding motif GxxG to GDDG) have a Δ*kphA*/*ΔeloR* phenotype (29), we wondered whether this was because KhpA^GDDG^ had reduced interaction with EloR. However, our BACTH assay showed that KhpA^GDDG^ successfully formed a complex with EloR (Fig. 4C), and we confirmed that *pbp2b* could be deleted in pneumococci expressing KhpA^GDDG^, as also reported by Zheng et al (29). This demonstrates that EloR interacts with KhpA because it fully depends on the ssRNA binding capacity of KhpA to form a functional EloR/KhpA complex.

### EloR recruits KhpA to the division site

KhpA and EloR have been shown to co-localize to the septal region of dividing cells (26, 29). Since they form heterodimers *in vivo*, we wondered if KhpA is recruited to mid-cell through its interaction with EloR. To explore this, the subcellular localization of sfGFP-fused KhpA was determined in wild type cells and in a Δ*eloR* mutant (Fig. 5). Mid-cell localization of KhpA-sfGFP was found in 75.4% of wild type cells, confirming previous findings (29). In contrast, KhpA-sfGFP was found at mid-cell in only 0.5% of the Δ*eloR* mutant cells. To show that it is the direct interaction between KhpA and EloR that localize KhpA to the division site and not some indirect effect of deleting the *eloR* gene, we fused sfGFP to the I61F and I61Y mutant versions of KhpA. As expected, KhpA^I61Y^-sfGFP, which does not bind EloR, lost its localization to mid-cell (found at mid-cell in only 2% of the cells). The monomeric KhpA^I61F^-sfGFP are still able to interact with EloR and displayed significantly higher degree of mid-cell localization (found at mid-cell in 19% of the cells). In accordance with these results expression of EloR^L239Y^, which cannot interact with KhpA, resulted in mislocalization of KhpA-sfGFP (Fig. 5). Together, these results strongly indicate that KhpA is recruited to mid-cell through complex formation with EloR.

**Fig. 5.**
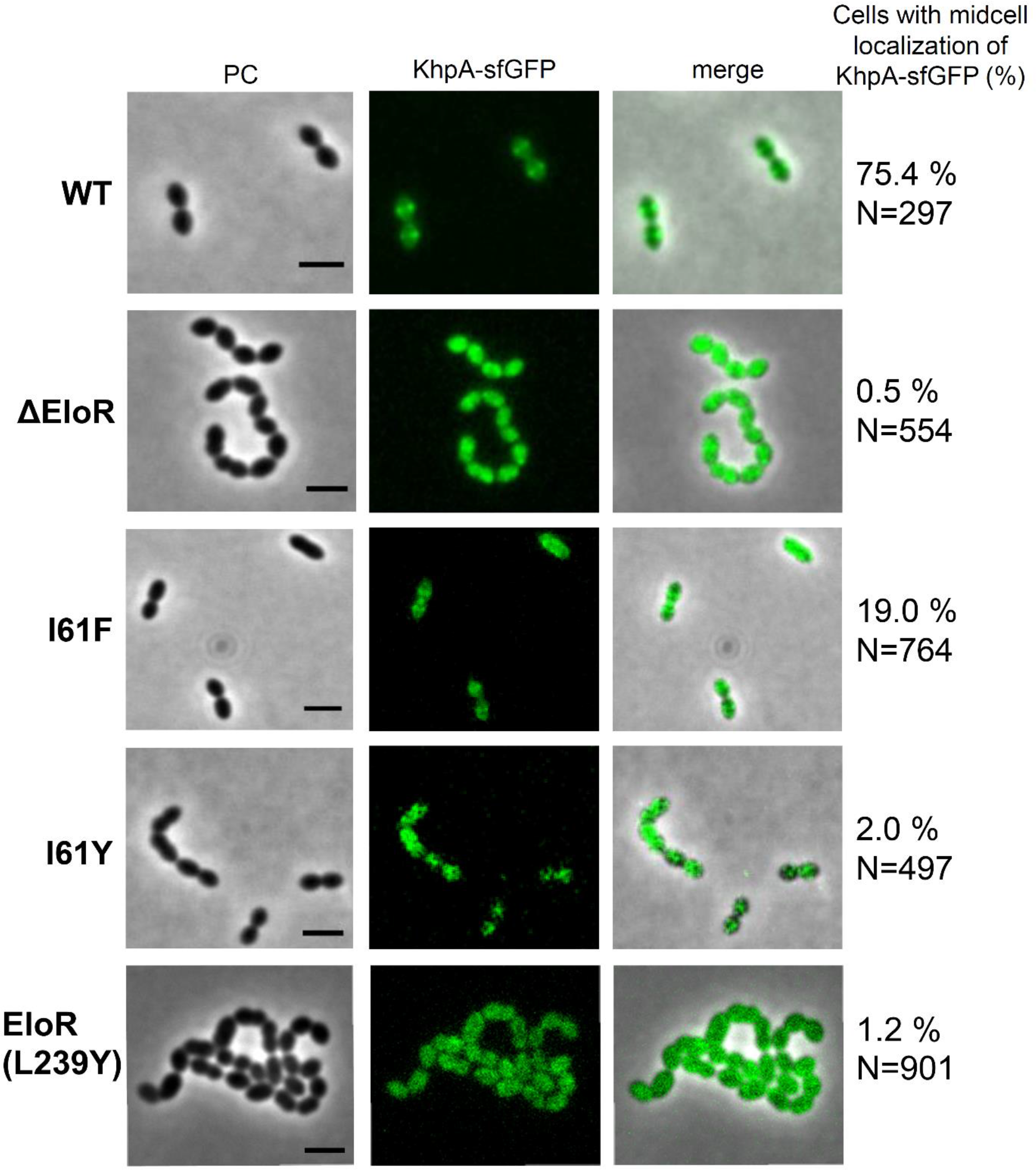
Micrographs showing the localization of KhpA-sfGFP in strain AW5 (wt), AW238 (Δ*eloR*), AW267 (KhpA^I61F^-sfGFP), AW321 (KhpA^I61Y^-sfGFP) and AW353 (EloR^L239Y^). The percent of cells having KhpA-sfGFP localized to mid-cell are indicated. Scale bars are 2 μm.

To determine whether the EloR/KhpA complex is recruited to the division zone during early, late or all stages of cell division, we compared the localization patterns of KhpA and FtsZ. FtsZ forms the division ring, which functions as a scaffold for a number of proteins found in the elongasome and divisome. FtsZ is therefore present at the division zone during initiation of new septa, cell elongation and cross wall synthesis, but it is not required for the final stage of daughter cell separation (12, 17). KhpA-sfGFP and FtsZ fused to the fluorescent marker mKate2 were co-expressed in *S. pneumoniae* (strain AW198), and fluorescence microscopy images demonstrate that mid-cell located KhpA-sfGFP follows the same localization pattern as FtsZ (Fig. 6). This shows that the EloR/KhpA complex is recruited to the division zone at the very early stage, and that it remains co-localized with the cell division machineries throughout the cell cycle. Note, however, that KhpA does not exclusively co-localized with FtsZ as it is also found throughout the cytoplasm.

**Fig. 6.**
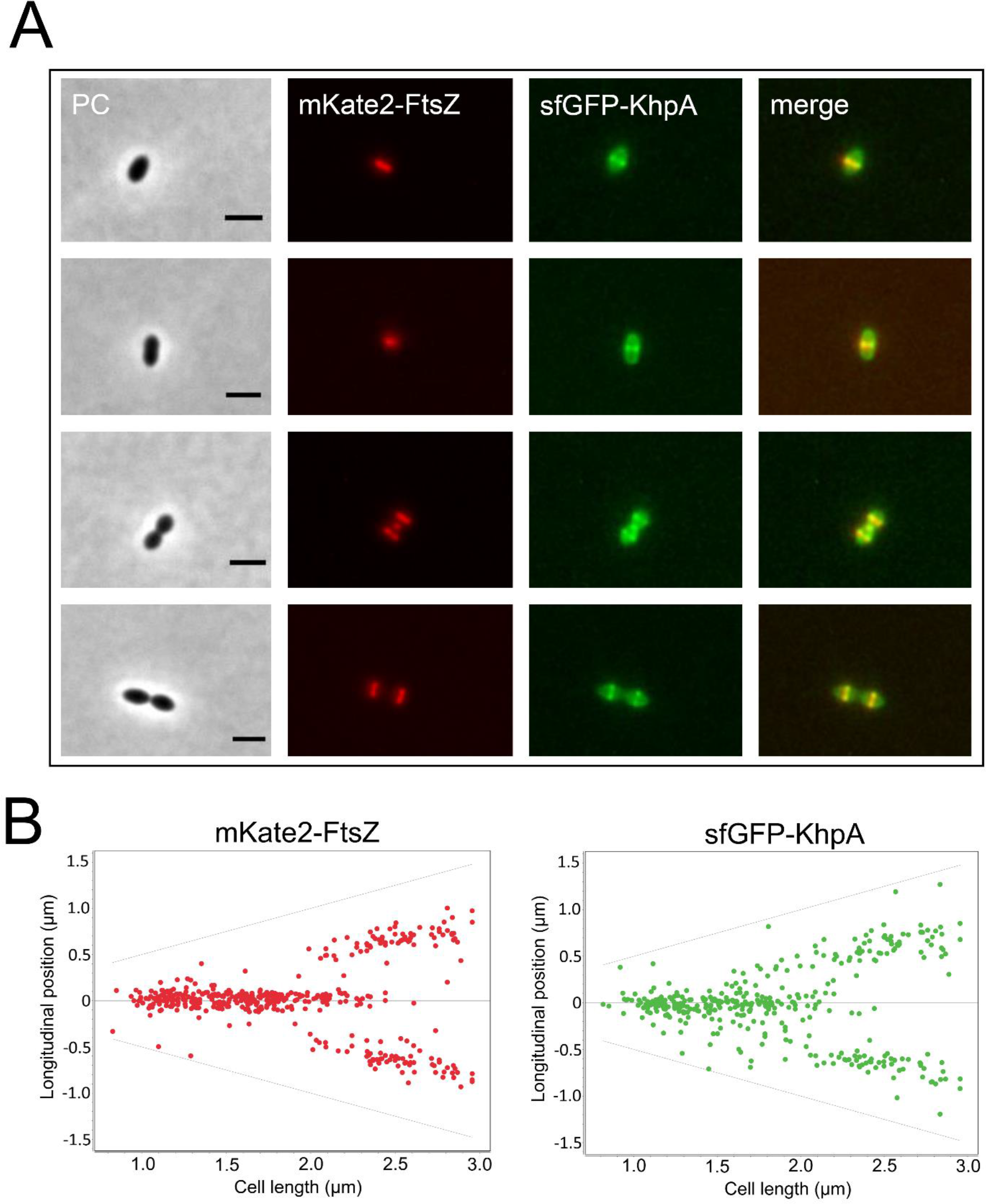
Localization of KhpA-sfGFP and mKate2-FtsZ at different stages of cell division. A. Microscopic examination of strain AW198 showed that KhpA-sfGFP co-localizes to the division site with FtsZ-mKate2 during cell division. Scale bars are 2 μm. B. The fluorescence maximum signals of FtsZ-mKate2 and KhpA-sfGFP plotted relative to cell length. 437 cells were analyzed.

## Discussion

It has been shown previously that Δ*khpA* and Δ*eloR* mutant strains are similar in several respects. They both exhibit less elongated cell morphologies, and are able to survive without PBP2b and other essential components of the elongasome (26, 29). The fact that Δ*khpA* and Δ*eloR* mutants have similar phenotypes could suggest that KhpA and EloR are acting at different steps in the same regulatory pathway. However, the finding that KhpA co-precipitates with EloR after formaldehyde crosslinking (29) suggests an alternative model, namely that they function as a single unit and that disruption of this complex gives rise to the phenotypes described above. The results presented in the present work prove that the latter model is correct. Disruption of the EloR/KhpA complex by introduction of site-specific amino acid substitutions, gives rise to shorter cells and renders the elongasome redundant. It is therefore likely that its role is to stimulate or control elongasome-mediated lateral cell wall synthesis. To do this, our results show that KhpA must be able to bind its target nucleic acid, which is most likely ssRNA. The typical binding surface of KH-domains can only accommodate four unpaired bases (28, 31), and consequently has low binding specificity. It is reasonable to assume that the RNA sequence motifs recognised by KhpA and the KH-II domain of EloR are different. Hence, by combining the two domains in a heterodimer the binding specificity and affinity for its target ssRNA(s) are substantially increased. The target RNA(s) bound by the EloR/KhpA complex might be ribosomal RNA, small noncoding RNA or mRNA. Identification of this RNA will be an important goal for future research seeking to understand the function of the EloR/KhpA system.

Our results show that KhpA also forms homodimers, which might have their own distinct biological function. The observed homomeric and heteromeric interactions of KhpA seem to be equally strong (see Fig. 3A), and it is therefore likely that both complexes forms *in vivo*. However, our preliminary studies did not detect any obvious functional deficits or major phenotypic changes associated with the KhpA^I61F^ mutation, i.e. the mutation disrupting the formation of KhpA homodimers without preventing the formation of EloR/KhpA heterodimers. As the KhpA monomers are arranged in an antiparallel orientation in the dimer, they will be able bind two successive sequence motifs on the same RNA strand. The binding of two motifs will increase the target sequence specificity considerably, and will make the RNA sequence motif recognized by the homodimer different from that recognized by the EloR/KhpA heterodimer. Considering this, and that the KhpA^I61F^ and KhpA^I61Y^ mutations give rise to completely different phenotypes, it is likely that the KhpA homodimers and EloR/KhpA heterodimers serve different biological functions.

The EloR/KhpA heterodimer contains three RNA-binding domains, i.e two domains from EloR (KH-II and R3H) and one from KhpA. The presence of several RNA-binding domains is a common feature of proteins containing KH-domains. As mentioned above, this increases target specificity and is also believed to have an important role in the folding of ssRNA sequences (31). Based on the present and previous studies (25, 26, 29), we know that the EloR/KhpA complex requires the combined action of all three RNA-binding domains to regulate cell elongation. However, it is not known whether all three domains bind to the same RNA strand, or if the KH-II^EloR^/KhpA complex binds one strand while the R3H domain binds another. The crystal structure of an EloR homolog from *Clostridium symbosium* (PDB 3GKU) suggests a dimeric structure (33), which in principle could bind two KhpA molecules resulting in a complex with a total of six RNA-binding domains. To test this possibility we used the BACTH system to determine if EloR from *S. pneumoniae* forms homodimers. The results were inconclusive as we obtained just a very weak positive signal (data not shown). Hence, we cannot conclude whether the biologically active complex between EloR and KhpA is dimeric (EloR/KhpA) or tetrameric (KhpA/EloR/EloR/KhpA).

Synthesis of the lateral cell wall takes place in an area close to the division septum, possibly where the division septum meets the periphery of the cell. Previous studies show that EloR and KhpA localize to the septal region (26, 29). Here, we show that KhpA homodimers are found throughout the cytoplasm (strain AW353) (Fig. 5), while KhpA/EloR heterodimers localize together with FtsZ to the division site (AW198) (Fig. 6). This finding support the notion that these homo- and heterodimers serve different functions. Since KhpA co-localizes with the FtsZ-ring throughout the cell cycle, it suggests that a functional EloR/KhpA complex is important during the stages of cell division, which involves active peptidoglycan synthesis, but not during the final stage of daughter cell separation. Of note, FtsZ has been reported to disappear from the septum prior to the essential divisome protein PBP2x (12). Since the EloR/KhpA complex closely follows the FtsZ localization pattern, it is compatible with the idea that the EloR/KhpA complex is involved in controlling the activity of the elongasome rather than controlling the divisome and septal cross-wall synthesis.

Zheng and co-workers report that the levels of FtsA, which together with FtsZ assembles into the division ring (6, 17, 34, 35), were elevated two-to threefold in Δ*eloR* and Δ*khpA* mutants. Their results suggest that EloR and KhpA bind 5’ untranslated regions of mRNAs, including the *ftsA* transcript, resulting in altered translation rates (29). In support of this hypothesis they found that *pbp2b* could be deleted in wild type cells overexpressing FtsA, although overexpression of FtsA could not fully restore the wild type phenotype of Δ*eloR*/Δ*khpA* cells (29). We attempted to reproduce the described effect of elevated FtsA levels in our D39 and R6 strains. However, despite using the exact same expression conditions, i.e. overexpression of *ftsA* and its 24 nt upstream region from a P_Zn_ zinc-inducible promoter, we were not successful. Nevertheless, translational control of specific mRNAs seems to be the most probable mode of action for the EloR/KhpA complex.

Interestingly, the *eloR* gene is co-transcribed with a gene called *yidC* in *S. pneumoniae* (36) and most likely in several other bacteria including *S. thermophilus*, *L. monocytogenes*, *B. subtilis*, *L. lactis*, *E. faecium* and *L. plantarum*. Such conserved co-transcription could indicate a functional relationship between the genes. YidC is an insertase that assists in co-translational insertion of membrane proteins into the lipid bilayer. It functions together with the SecYEG translocon, the signal recognition particle (SRP) and the SRP-receptor FtsY. During co-translational protein targeting to the SecYEG translocon, the SRP-ribosome-nascent protein chain complex is first targeted to FtsY, which delivers the chain to the SecYEG translocon channel. The function of YidC is to facilitate the release of the transmembrane domains of inner membrane proteins from the channel into the lipid bilayer (37, 38). Having this in mind, it is tempting to speculate that the EloR/KhpA complex could be involved in regulating the expression and insertion of specific membrane proteins involved in cell elongation through translational control.

## Materials and Methods

### Bacterial strains, cultivation and transformation

All strains used in this work are listed in Table 1. *E. coli* strains were grown in LB broth at 37°C with shaking (200 rpm), or on LB plates at 37°C unless otherwise indicated. When necessary the following antibiotics were used: kanamycin (50 μg/ml) and ampicillin (100 μg/ml). Transformation experiments were performed with chemically competent cells using the heat shock method at 42°C for 45 seconds. *S. pneumoniae* were grown in C medium (39) or on Todd Hewitt-agar plates at 37°C. Agar plates were incubated in anaerobic chambers using AnaeroGenTM bags from Oxoid. When necessary, kanamycin (400 μg/ml) and streptomycin (200 μg/ml) were employed for selection of transformants. In order to knock out genes or introduce mutations, natural genetic transformation was employed. For transformation experiments, the culture was grown to an OD_550_ of 0.05-0.1 and mixed with the transforming DNA (100-200 ng) and CSP1, which was added to a final concentration of 250 ng/ml. After 2 hours of incubation at 37°C, 30 μl of the culture was plated on TH-agar containing the appropriate antibiotic followed by incubation at 37°C over night. To investigate growth rates of different mutants, cultures were grown to an OD_550_ of 0.2, diluted to OD_550_ = 0.05, and grown in 96-well Corning NBS clear-bottom plates in a Synergy H1 Hybrid Reader (BioTek). The OD_550_ was measured automatically every 5 minutes for 20 hours.

**Table 1.** *S. pneumoniae* strains used in the present study.

| Name | Relevant characteristics | Reference |
| --- | --- | --- |
| R704 | R6 derivative, *comA::ermAM*; Ery<sup>R</sup> | JP. Claverys* |
| RH425 | R704, but streptomycin resistant; Ery<sup>R</sup>, Sm<sup>R</sup> | (49) |
| DS420 | $\Delta comA$, $\Delta khpA$; Ery<sup>R</sup>, Sm<sup>R</sup> | This work |
| DS428 | $\Delta comA$, $\Delta khpA$, $\Delta pbp2b::janus$; Ery<sup>R</sup>, Kan<sup>R</sup> | This work |
| AW5 | $\Delta comA$, *khpA-sfgfp*; Ery<sup>R</sup>, Sm<sup>R</sup> | This work |
| AW24 | $\Delta comA$, *khpA<sup>GDDG</sup>*; Ery<sup>R</sup>, Sm<sup>R</sup> | This work |
| AW27 | $\Delta comA$, *khpA<sup>GDDG</sup>*, $\Delta pbp2b::janus$; Ery<sup>R</sup>, Kan<sup>R</sup> | This work |
| AW198 | $\Delta comA$, *khpA-sfgfp*, *ftsZ-mKate2-Km*; Ery<sup>R</sup>, Km<sup>R</sup>, Sm<sup>R</sup> | This work |
| AW212 | $\Delta comA$, *khpA<sup>I61F</sup>*; Ery<sup>R</sup>, Sm<sup>R</sup> | This work |
| AW238 | $\Delta comA$, *khpA-sfgfp*, $\Delta eloR$; Ery<sup>R</sup>, Sm<sup>R</sup> | This work |
| AW267 | $\Delta comA$, *khpA<sup>I61F</sup>-sfgfp*; Ery<sup>R</sup>, Sm<sup>R</sup> | This work |
| AW275 | $\Delta comA$, *khpA<sup>I61Y</sup>*; Ery<sup>R</sup>, Sm<sup>R</sup> | This work |
| AW279 | $\Delta comA$, *eloR<sup>L239Y</sup>*; Ery<sup>R</sup>, Sm<sup>R</sup> | This work |
| AW313 | $\Delta comA$, *khpA<sup>I61Y</sup>*, $\Delta pbp2b::janus$; Ery<sup>R</sup>, Kan<sup>R</sup> | This work |
| AW314 | $\Delta comA$, *eloR<sup>L239Y</sup>*, $\Delta pbp2b::janus$; Ery<sup>R</sup>, Kan<sup>R</sup> | This work |
| AW321 | $\Delta comA$, *khpA<sup>I61Y</sup>-sfgfp*; Ery<sup>R</sup>, Sm<sup>R</sup> | This work |
| AW334 | $\Delta comA$, *flag-eloR<sup>L239C</sup>*; Ery<sup>R</sup>, Sm<sup>R</sup> | This work |
| AW336 | $\Delta comA$, *flag-eloR<sup>L239C</sup>*, *khpA<sup>I61C</sup>*; Ery<sup>R</sup>, Sm<sup>R</sup> | This work |
| AW353 | $\Delta comA$, *khpA-sfgfp*, *eloR<sup>L239Y</sup>*; Ery<sup>R</sup>, Sm<sup>R</sup> | This work |
| SPH446 | $\Delta comA$, $\Delta eloR$, $\Delta pbp2b::janus$; Ery<sup>R</sup>, Kan<sup>R</sup> | (26) |
| SPH448 | $\Delta comA$, *flag-eloR*; Ery<sup>R</sup>, Sm<sup>R</sup> | (26) |
| RR66 | D39 derivative, *ftsZ-mKate2*, Kan<sup>R</sup> | (42) |

\*Gift from Professor Jean-Pierre Claverys, CNRS, Toulouse, France.

### Construction of genetic mutants, gene fusions and point mutations

DNA amplicons used in transformation experiments were created with overlap extension PCR as previously described (40). Genes were knocked out using a Janus cassette (41). The cassettes were created with sequences of ~1000 bp homologous to the flanking sequences of the insertion site in the genome. The same technique was employed when introducing point mutations or fusion genes. Primers used to create these amplicons are listed in Table S1. The ftsZ-mKate2 fusion gene together with a kanamycin resistance cassette was amplified from genomic DNA of strain RR66 (42). All constructs were verified with PCR and Sanger Sequencing.

### SDS-PAGE and immunoblotting

The strain RH425, SPH448, AW334 and AW336 were grown to an OD_550_ of 0.3 in a culture volume of 45 ml. The cells were harvested at 4000 x *g*, and resuspended in 200 μl 1 x SDS sample buffer not containing any reducing agents. The samples were then split in two, and β-mercaptoethanol was added to one parallel half of the samples to a final concentration of 100 mM. All the samples (including the non-reduced) were heated at 100 °C for 10 minutes. The cell lysates were separated on a 15 % polyacrylamide gel with buffer conditions as previously described (43). For immunodetection purposes, the separated proteins were electroblotted onto a PVDF membrane (BioRad), and flag-EloR was detected with α-flag antibodies as previously described (44).

### BACTH-assay

The bacterial adenylate cyclase two hybrid (BACTH) assay, is based on the functional complementation of T18 and T25, two domains of the *B. pertussis* adenylate cyclase (CyaA) (30). When these domains are brought in close proximity to each other, they can actively produce cAMP. The production of cAMP leads to activation of the catabolite activator protein CAP, which in a complex with cAMP activates expression of a reporter gene placed behind the cAMP/CAP promoter. The reporter gene used in this system encodes the β-galactosidase enzyme. In order to investigate the interaction between two proteins, we cloned genes encoding the proteins of interest in frame with either the T25 -or the T18-encoding sequences in plasmids provided by the manufacturer (Euromedex). The plasmids used in this study are listed in Table S2. Next, two plasmids, each expressing one protein fused to either T18 or T25 were transformed into *E. coli* BTH101 cells (a *cya*^-^ strain). After overnight incubation on LB plates containing kanamycin (50 μg/ml) and ampicillin (100 μg/ml), five colonies from each transformation were grown in LB containing the appropriate antibiotics. When reaching an OD^600^ of 0.2, three μl of the cell cultures were spotted onto LB plates containing 0.5 mM IPTG (to induce expression of the fusion genes), X-gal (40 μg/ml), kanamycin (50 μg/ml) and ampicillin (100 μg/ml). After an overnight incubation at 30°C, results were interpreted as positive or negative based on the color of the spot. A positive interaction between the proteins of interest will result in blue spots on a plate. In addition, the production of β-galactosidase reporter was measured quantitatively by performing β-galactosidase assays using ortho-nitrophenyl-β-galactoside (ONPG) as substrate. *E. coli* BTH101 containing plasmids with T18 and T25-fused genes were grown in the presence of kanamycin (50 μg/ml) and ampicillin (100 μg/ml) to OD_600_ = 0.4-0.5. Then the cells were diluted to OD_600_ = 0.05 in similar medium also containing 0.5 mM IPTG. The cells were incubated at 30 °C with shaking for 4 hours. Cells from one ml culture were lysed using 0.5 g of ≤106 μm glass beads (Sigma) and bead beating at 6.5 m/s for 3x20 seconds. Then the β-galactosidase activity in 100 μl cell lysate was determined following the protocol of Steinmoen *et al.* (45).

### Microscopy and cell shape distribution analyses

The subcellular localization of different point mutated versions of the KhpA proteins was examined by fluorescence microscopy. The mutated proteins in question were fused to sfGFP (42) via a short glycine-linker (GGGGG). sfGFP fusions were expressed in the native *khpA* locus in the *S. pneumoniae* genome (strains AW5, AW198, AW238, AW267, AW321 and AW353).

The cell morphology and cell shape distributions were examined by phase contrast microscopy. Microscopy experiments were performed by growing the strains to an OD_550_ of 0.1 before immobilizing the cells on a microscopy slide using 1.2 % low melting agarose (Biorad) in PBS. Phase contrast images and GFP fluorescence images were obtained using a Zeiss AxioObserver with ZEN Blue software, and an ORCA-Flash 4.0 V2 Digital CMOS camera (Hamamatsu Photonics) using a 1003 phase-contrast objective. The ImageJ plugin MicrobeJ (46) was used to analyze the cell shape and the subcellular localization of KhpA-sfGFP and FtsZ-mKate2. Cells were segmented using the phase contrast images. Cell shape distributions were made by calculating length/width for the individual cell and the significance of the differences between distributions were determined using a two-sample t-test. To determine the percentage of cells with mid-cell localized KhpA-sfGFP, the GFP fluorescence profiles were plotted for the individual cells. KhpA-sfGFP was scored as mid-cell localized when a fluorescence maximum peak was found in the mid-cell area (between 40-60 % of the cell length), and the percentage of cells with mid-cell localized KhpA-sfGFP was calculated. To analyze the subcellular localization of FtsZ-mKate2 and KhpA-sfGFP, the Maxima-option in MicrobeJ was used.

### 3D-modelling

The online structure determination tool iTasser was used to predict the 3D-structure of KhpA. It uses algorithms to predict protein 3D structure based on the amino acid sequence and known, published structures (47). The ZDOCK server was used to predict the interaction surface in a KhpA homodimer (48). Based on the predicted interaction surface in a KhpA homodimer, we created point mutated versions of KhpA, introduced these into the BACTH system, and tested interactions between mutated KhpA proteins and between mutated KhpA and wild type EloR.

## Acknowledgements

This work was partly funded by a grant given by the Research Council of Norway. The authors have no conflict of interest with regard to the data presented in this study.

## References

1. Vollmer W, Blanot D, & de Pedro MA (2008) Peptidoglycan structure and architecture. FEMS Microbiol Rev 32(2):149–167.

2. Dramsi S, Magnet S, Davison S, & Arthur M (2008) Covalent attachment of proteins to peptidoglycan. FEMS Microbiol Rev 32(2):307–320.

3. Brown S, Santa Maria JP, Jr., & Walker S (2013) Wall teichoic acids of Gram-positive bacteria. Annu Rev Microbiol 67:313–336.

4. Bazaka K, Crawford RJ, Nazarenko EL, & Ivanova EP (2011) Bacterial extracellular polysaccharides. Adv Exp Med Biol 715:213–226.

5. Sørensen UB, Henrichsen J, Chen HC, & Szu SC (1990) Covalent linkage between the capsular polysaccharide and the cell wall peptidoglycan of Streptococcus pneumoniae revealed by immunochemical methods. Microb Pathog 8(5):325–334.

6. Pinho MG, Kjos M, & Veening JW (2013) How to get (a)round: mechanisms controlling growth and division of coccoid bacteria. Nature reviews. Microbiology 11(9):601–614.

7. Zapun A, Vernet T, & Pinho MG (2008) The different shapes of cocci. FEMS Microbiol Rev 32(2):345–360.

8. Cho H, et al. (2016) Bacterial cell wall biogenesis is mediated by SEDS and PBP polymerase families functioning semi-autonomously. Nat Microbiol:16172.

9. Emami K, et al. (2017) RodA as the missing glycosyltransferase in Bacillus subtilis and antibiotic discovery for the peptidoglycan polymerase pathway. Nat Microbiol 2:16253.

10. Sauvage E, Kerff F, Terrak M, Ayala JA, & Charlier P (2008) The penicillin-binding proteins: structure and role in peptidoglycan biosynthesis. FEMS Microbiol Rev 32(2):234–258.

11. Berg KH, Stamsås GA, Straume D, & Håvarstein LS (2013) Effects of low PBP2b levels on cell morphology and peptidoglycan composition in Streptococcus pneumoniae R6. J Bacteriol 195(19):4342–4354.

12. Tsui HC, et al. (2014) Pbp2x localizes separately from Pbp2b and other peptidoglycan synthesis proteins during later stages of cell division of Streptococcus pneumoniae D39. Mol Microbiol 94(1):21–40.

13. Land AD, et al. (2013) Requirement of essential Pbp2x and GpsB for septal ring closure in Streptococcus pneumoniae D39. Mol Microbiol 90(5):939–955.

14. Perez-Nunez D, et al. (2011) A new morphogenesis pathway in bacteria: unbalanced activity of cell wall synthesis machineries leads to coccus-to-rod transition and filamentation in ovococci. Mol Microbiol 79(3):759–771.

15. Wheeler R, Mesnage S, Boneca IG, Hobbs JK, & Foster SJ (2011) Super-resolution microscopy reveals cell wall dynamics and peptidoglycan architecture in ovococcal bacteria. Mol Microbiol 82(5):1096–1109.

16. Garner EC, et al. (2011) Coupled, circumferential motions of the cell wall synthesis machinery and MreB filaments in B. subtilis. Science 333(6039):222–225.

17. Mura A, et al. (2016) Roles of the essential protein FtsA in cell growth and division in Streptococcus pneumoniae. J Bacteriol.

18. Jacq M, et al. (2015) Remodeling of the Z-Ring Nanostructure during the Streptococcus pneumoniae Cell Cycle Revealed by Photoactivated Localization Microscopy. MBio 6(4).

19. Straume D, Stamsås GA, Berg KH, Salehian Z, & Håvarstein LS (2017) Identification of pneumococcal proteins that are functionally linked to penicillin-binding protein 2b (PBP2b). Mol Microbiol 103(1):99–116.

20. Fleurie A, et al. (2012) Mutational dissection of the S/T-kinase StkP reveals crucial roles in cell division of Streptococcus pneumoniae. Mol Microbiol 83(4):746–758.

21. Novakova L, et al. (2010) Identification of multiple substrates of the StkP Ser/Thr protein kinase in Streptococcus pneumoniae. J Bacteriol 192(14):3629–3638.

22. Beilharz K, et al. (2012) Control of cell division in Streptococcus pneumoniae by the conserved Ser/Thr protein kinase StkP. Proc Natl Acad Sci U S A 109(15):E905–913.

23. Zucchini L, et al. (2018) PASTA repeats of the protein kinase StkP interconnect cell constriction and separation of Streptococcus pneumoniae. Nat Microbiol 3(2):197–209.

24. Sun X, et al. (2010) Phosphoproteomic analysis reveals the multiple roles of phosphorylation in pathogenic bacterium Streptococcus pneumoniae. J Proteome Res 9(1):275–282.

25. Ulrych A, et al. (2016) Characterization of pneumococcal Ser/Thr protein phosphatase phpP mutant and identification of a novel PhpP substrate, putative RNA binding protein Jag. BMC Microbiol 16(1):247.

26. Stamsås GA, et al. (2017) Identification of EloR (Spr1851) as a regulator of cell elongation in Streptococcus pneumoniae. Mol Microbiol 105(6):954–967.

27. Grishin NV (1998) The R3H motif: a domain that binds single-stranded nucleic acids. Trends Biochem Sci 23(9):329–330.

28. Valverde R, Edwards L, & Regan L (2008) Structure and function of KH domains. FEBS J 275(11):2712–2726.

29. Zheng JJ, Perez AJ, Tsui HT, Massidda O, & Winkler ME (2017) Absence of the KhpA and KhpB (JAG/EloR) RNA-binding proteins suppresses the requirement for PBP2b by overproduction of FtsA in Streptococcus pneumoniae D39. Mol Microbiol 106(5):793–814.

30. Karimova G, Pidoux J, Ullmann A, & Ladant D (1998) A bacterial two-hybrid system based on a reconstituted signal transduction pathway. Proceedings of the National Academy of Sciences 95(10):5752–5756.

31. Nicastro G, Taylor IA, & Ramos A (2015) KH-RNA interactions: back in the groove. Curr Opin Struct Biol 30:63–70.

32. Hollingworth D, et al. (2012) KH domains with impaired nucleic acid binding as a tool for functional analysis. Nucleic acids research 40(14):6873–6886.

33. Tan K, Keigher L, Jedrzejczak R, Babnigg G, & Joachimiak A (The crystal structure of a probable RNA-binding protein from Clostridium symbiosum ATCC 14940). (http://www.rcsb.org/structure/3GKU).

34. Bisson-Filho AW, et al. (2017) Treadmilling by FtsZ filaments drives peptidoglycan synthesis and bacterial cell division. Science 355(6326):739–743.

35. Yang X, et al. (2017) GTPase activity-coupled treadmilling of the bacterial tubulin FtsZ organizes septal cell wall synthesis. Science 355(6326):744–747.

36. Slager J, Aprianto R, & Veening JW (2018) Deep genome annotation of the opportunistic human pathogen Streptococcus pneumoniae D39. Nucleic Acids Res https://doi.org/10.1093/nar/gky725.

37. Wu ZC, de Keyzer J, Berrelkamp-Lahpor GA, & Driessen AJ (2013) Interaction of Streptococcus mutans YidC1 and YidC2 with translating and nontranslating ribosomes. J Bacteriol 195(19):4545–4551.

38. Steinberg R, Knupffer L, Origi A, Asti R, & Koch HG (2018) Co-translational protein targeting in bacteria. FEMS Microbiol Lett 365(11).

39. Lacks S & Hotchkiss RD (1960) A study of the genetic material determining an enzyme activity in pneumococcus. Biochimica et biophysica acta 39(3):508–518.

40. Higuchi R, Krummel B, & Saiki R (1988) A general method of in vitro preparation and specific mutagenesis of DNA fragments: study of protein and DNA interactions. Nucleic acids research 16(15):7351–7367.

41. Sung C, Li H, Claverys J, & Morrison D (2001) An rpsL cassette, janus, for gene replacement through negative selection in Streptococcus pneumoniae. Applied and environmental microbiology 67(11):5190–5196.

42. van Raaphorst R, Kjos M, & Veening JW (2017) Chromosome segregation drives division site selection in Streptococcus pneumoniae. Proc Natl Acad Sci U S A 114(29):E5959–E5968.

43. Laemmli UK (1970) Cleavage of structural proteins during the assembly of the head of bacteriophage T4. nature 227(5259):680.

44. Stamsås GA, Straume D, Salehian Z, & Håvarstein LS (2017) Evidence that pneumococcal WalK is regulated by StkP through protein-protein interaction. Microbiology 163(3):383–399.

45. Steinmoen H, Knutsen E, & Håvarstein LS (2002) Induction of natural competence in Streptococcus pneumoniae triggers lysis and DNA release from a subfraction of the cell population. Proceedings of the National Academy of Sciences 99(11):7681–7686.

46. Ducret A, Quardokus EM, & Brun YV (2016) MicrobeJ, a tool for high throughput bacterial cell detection and quantitative analysis. Nature microbiology 1(7):16077.

47. Roy A, Kucukural A, & Zhang Y (2010) I-TASSER: a unified platform for automated protein structure and function prediction. Nature protocols 5(4):725.

48. Pierce BG, et al. (2014) ZDOCK server: interactive docking prediction of protein-protein complexes and symmetric multimers. Bioinformatics 30(12):1771–1773.

49. Johnsborg O & Håvarstein LS (2009) Pneumococcal LytR, a protein from the LytR-CpsA-Psr family, is essential for normal septum formation in Streptococcus pneumoniae. J Bacteriol 191(18):5859–5864.

